# Adaptation of a Mutual Exclusivity Framework to Identify Driver Mutations within Biological Pathways

**DOI:** 10.1101/2023.09.19.558469

**Authors:** Xinjun Wang, Caroline Kostrzewa, Allison Reiner, Ronglai Shen, Colin Begg

## Abstract

Distinguishing genomic alterations in cancer genes that have functional impact on tumor growth and disease progression from the ones that are passengers and confer no fitness advantage has important clinical implications. Evidence-based methods for nominating drivers are limited by existing knowledge on the oncogenic effects and therapeutic benefits of specific variants from clinical trials or experimental settings. As clinical sequencing becomes a mainstay of patient care, applying computational methods to mine the rapidly growing clinical genomic data holds promise in uncovering novel functional candidates beyond the existing knowledge-base and expanding the patient population that could potentially benefit from genetically targeted therapies. We propose a statistical and computational method (MAGPIE) that builds on a likelihood approach leveraging the mutual exclusivity pattern within an oncogenic pathway for identifying probabilistically both the specific genes within a pathway and the individual mutations within such genes that are truly the drivers. Alterations in a cancer gene are assumed to be a mixture of driver and passenger mutations with the passenger rates modeled in relationship to tumor mutational burden. A limited memory BFGS algorithm is used to facilitate large scale optimization. We use simulations to study the operating characteristics of the method and assess false positive and false negative rates in driver nomination. When applied to a large study of primary melanomas the method accurately identified the known driver genes within the RTK-RAS pathway and nominated a number of rare variants with previously unknown biological and clinical relevance as prime candidates for functional validation.

## Introduction

It is now well known that cancer is a genetic disease that develops through the accumulation of somatic mutations. When individual tumors are subjected to mutation analysis countless mutations are identified. A major challenge is to identify so-called driver mutations, the ones that are pivotal in producing the uncontrolled tumor growth. A major technical concept that has influenced research in this field is mutual exclusivity. If somatic mutations in two (or more) genes tend not to occur together in the same tumor then this is evidence that the disruption of the genes involved are leading to similar effects. Presence of such mutual exclusivity is strong evidence that the genes are cancer genes. Important findings on mutual exclusivity through existing large-scale sequencing studies include *EGFR* and *KRAS* in lung adenocarcinoma^1, 2, 3^, and the RTK-RAS pathway primarily involving *BRAF, NRAS*, and *NF1* in melanoma^4, 5^.

Studies of this phenomenon usually involve searching for evidence of mutual exclusivity of genes in a “pathway” of genes that are believed to possess related effects. A number of authors have studied this problem from a statistical perspective, developing several techniques. The methods called Dendrix^6^ and Mutex^7^ define novel criteria or score metrics based on which they search for the mutually exclusive gene sets using a greedy approach. MEMo^8^ and gcMECM^9^ are based on a search for mutually exclusive genes using graph or network-based approaches. WeSME^10^ and FaME^11^ employ computational-oriented methods that can scale up to genome-wide analysis. CoMEt^12^ and WExT^13^ employ a permutation-based test for mutual exclusivity. A method by Szczurek et al.^14^, MEGSA^15^, DISCOVER^16^, TiMEx^17^ and MEScan^18^ utilize probabilistic model-based tests to assess the significance of mutual exclusivity in a given gene set.

Most of these methods focus on de novo search of gene sets (from the over 22,000 genes in the human genome) that display mutual exclusivity. In this article we turn our attention to leveraging the mathematical property of mutual exclusivity for identifying probabilistically both the specific genes within a pathway and the individual mutations within such genes that are truly the drivers. The evaluation of mutual exclusivity will occur in pre-defined pathways of genes, i.e., collections of genes that have been shown in previous research to share biological functions^19^. A major assumption is that, within any given pathway, only one of the mutations observed is a “driver” for that tumor, though there could be additional drivers in other pathways.

We build our method around the likelihood function developed by Hua et al^15^. In their model a mutation in a given gene can represent either a driver or a passenger mutation. Passenger mutations in this context represent random, non-consequential background somatic mutations that are a result of the genetic instability commonplace in tumor cells. The global patterns of mutual exclusivity in the data allow this model to identify the proportion of tumors that possess a driver in the pathway, based on the assumption that the driver mutations are completely mutually exclusive, i.e., a tumor can contain at most 1 driver in the pathway under investigation. A test of the hypothesis that this proportion is zero thus represents a test of mutual exclusivity of the set of genes under consideration. By applying this repeatedly to different subsets of genes in the pathway one can find the most significant subset, and thus conclude that this subset of genes consists of the drivers. Hua et al. made the assumption that the relative proportions of drivers versus passengers in each gene have a common proportionality. In our approach we relax this assumption, allowing us to estimate the proportions of driver mutations for each gene and then, through Bayes’ rule, to determine probabilistically which mutations in each tumor are drivers and which ones are passengers. By mapping all of these probabilities we can determine for which genes drivers predominate. Also, these individual probabilities allow us to shed light on which types of mutation within any given gene show up as drivers frequently. These results are driven fundamentally by the global empirical patterns of mutual exclusivity in the dataset.

In summary we show that our method goes beyond using statistical tests for mutual exclusivity to create a framework for inferring probabilistically which genes have the strongest evidence as drivers, and which mutations within these genes are specifically identified as drivers. We show through detailed analysis of the RTK-RAS pathway in a large sample of melanomas how the method confirms the prominence of a small number of recurring mutations in well-known genes in this pathway and also identifies some rare individual mutations that appear to be important. Our method has been implemented in a Python package named MAGPIE (<u>m</u>utual exclusivity <u>a</u>nalysis of cancer <u>g</u>enes and variants and their <u>p</u>robability of be<u>i</u>ng driv<u>e</u>r).

## Methods

Our data framework involves a set of *N* tumors and the analysis is restricted to a set of *M* genes in the pathway under consideration. That is, the analytic framework is pathway-specific. The data could include sequencing of genes in other pathways, but analyses of these other pathways would be conducted independently. That is, any given tumor may have multiple driver genes, but a key assumption is that there can only be one driver mutation in the pathway under consideration.

### MEGSA Framework

We initially construct our strategy on the MEGSA (mutually exclusive gene set analysis) likelihood-based analysis introduced by Hua et al^15^. Let ***x***_***i***_ =(*x*_*i*1_, …,*x*_*iM*_) denote the observed binary mutation status of tumor *i*(*i* = 1, …,*N*), where *x*_*i j*_ = 1 if an alteration is observed in the *j*^*th*^ gene (*j* =1, …,*M*) and 0 otherwise. Any observed mutation must be either a driver mutation or a passenger mutation. Only one driver mutation from the pathway is possible in a given tumor. As a result, all driver mutations observed must be mutually exclusive, i.e., no two drivers can occur in a given tumor. We define *γ*(0 ≤ *γ* ≤ 1) as the proportion of tumors in the study cohort that have a driver mutation in the pathway under investigation. Let *p*_*j*_ denote the relative frequency of such tumors that possess a driver mutation in the *j*^*th*^ gene in the pathway with 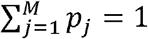 and 0 ≤ *p*_*j*_ ≤ 1 Independent of driver mutations, each gene has a constant passenger mutation rate, denoted by *π*_*j*_ for the *j*^*th*^ gene. The log likelihood of the observed data is

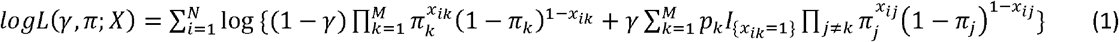

For the purpose of developing a statistical test of mutual exclusivity, Hua et al. made the assumption that the relative frequencies of driver mutations in each gene are proportional to the passenger mutation rates, i.e., *p*_*j*_ ∝ *π*_*j*_. Thus, the log likelihood is reduced to

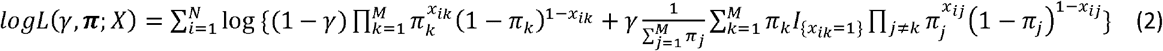

To test the fundamental null hypothesis that there is no mutual exclusivity in the pathway a likelihood ratio test can be employed. This is in effect a test of the hypothesis *H*_0_: *γ* = 0 versus the alternative (*H*_1_) that *γ* >0, where the test statistic has a null distribution of 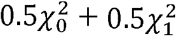.^15^

### Proposed Approach

MEGSA is a powerful tool to quantify the overall mutual exclusivity in the pathway or to select a subset of genes that reflect mutual exclusivity most strongly. Although the new method we propose in this article is adapted from MEGSA framework, the problem it solves is fundamentally different. Our method is designed to identify specifically which tumors possess a driver mutation and to identify the driver if more than one mutation in the pathway is present. It is further able to identify which variants within a given gene have the capacity to be driver variants.

We follow the model proposed in equation (1). However, unlike Hua et al. we do not assume *p*_*j*_ *∝ π*_*j*_ a critical assumption in the MEGSA approach. We further reformulate the likelihood into a mixture model framework. Assume that the gene membership of the driver mutations for tumor *i* in the study cohort is denoted by ***z***_***i***_ = (*z*_*i*0_, *z*_*i*1_, …,*z*_*iM*_). We emphasize that ***z***_***i***_ is unobserved and must be inferred. Tumors with *z*_*ij*_ =1,*j* > 0 have driver mutations in the *j*^*th*^ gene. Tumors with *z*_*i* 0_ = 1 do not possess a driver mutation. Let ***τ*** = (*τ*_0,_ *τ*_1_, …, *τ*_*M*_) denote the vector of proportions of tumors having each gene-specific driver mutation, i.e., *τ*_*j*_ *= p*(*z*_*ij*_ = 1). Note that in this new notation *τ*_*j*_ represents the absolute relative frequency of tumors with drivers in the *j*^*th*^ gene, while in the earlier notation *p*_*j*_ represents the corresponding relative frequency of the presence of a driver in the *j*^*th*^ gene among tumors that have drivers in the pathway. Following a standard mixture model framework, the log likelihood of the observed data is

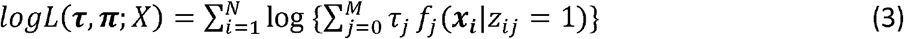

where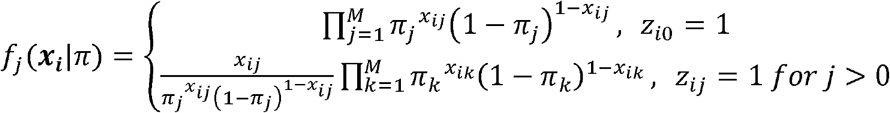 is the cluster-specific probability density function of ***x***_***i***_.

Equation (3) and equation (1) are in fact equivalent. To be specific, *τ*_0_ =1 − *γ* and *τ*_*j*_*= γp*_*j*_ for *j* >0. As before, *γ* = 1− *τ*_0_ quantifies the overall influence of a pathway, i.e., the proportion of tumors with a driver mutation in the pathway, while *τ*_*j*_,*j* >0, quantifies the relative frequency for which gene *j* is the driver. One of the advantages of using a mixture model framework is that *τ*_*j*_ ^*’*^*s* are defined both under the null hypothesis of no driver mutations in the cohort (i.e., *τ*_0_ = 1 or *γ* = 0) and under the alternative (i.e., *τ*_0_ <1 or *γ* > 0 :there exists evidence of a mutually exclusive pattern), while for the MEGSA model the *p*_*j*_^’^*s* are undefined under the null hypothesis. For the rest of this article, we follow the mixture model framework.

Parameters {*τ*_*j*_} and {*π*_*j*_}can be estimated using the EM algorithm^20^. The complete data log likelihood is

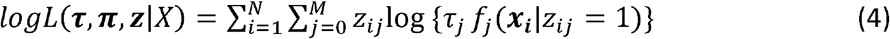

In the E-step, we compute the posterior probability *w*_*ij*_ = *p*(*z*_*ij*_=1|***x***_***i***_)at the *t*^*th*^ iteration:

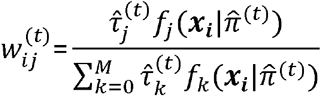

In the M-step, equation (4) is maximized in terms of {*τ*_*j*_}and { *π*_*j*_} with *w*_*ij*_ fixed at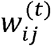:

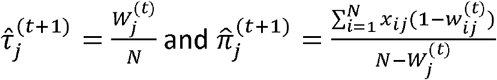

where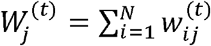.

In general, given initial estimates 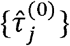 and 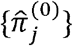, the EM algorithm then iterates between E-step and M-step until the estimates converge.

### Adjustment for Tumor Mutational Burden

Up to now the model has been based on the assumption that the passenger mutation rates {*π*_*j*_}are considered constant across the set of tumors. In fact, this assumption is quite unrealistic since the overall tumor mutational burden is known to vary widely across tumors and is, in many oncologic settings, an influential prognostic factor^21^. To address this important potential confounder, we extend the method to allow adjustment for the effect of mutational burden in the model. Let *y*_*i*_ denote the tumor mutational burden for tumor *i*. This represents the overall propensity for mutations to occur in a specific tumor. Due to this dependency, we now identify the passenger mutation rates using {*π*_*i j*_}rather than {*π*_*j*_}. In our later example mutational burden is represented by the total number of mutations observed across all genes that are genotyped, not just those in the pathway under investigation. Since *π*_*ij*_ is bounded by 0 and 1, a natural approach to adjust for mutational burden is to use

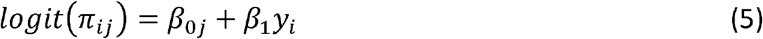

where *β*_0*j*_ represents the baseline log odds of the passenger mutation rate for the *j* ^*th*^ gene, and *β*_1_ measures the common influence of mutational burden on the passenger mutation rate for all genes in the pathway. Since we are primarily interested in estimating the probabilities {*τ*_*k*_}representing which gene is the driver, *β*_0*j*_ and *β*_1_ are effectively nuisance parameters in the model. The conditional data density for ***x***_***i***_ | *y*_*i*_ is

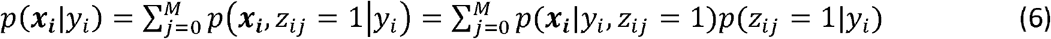

We assume **z**_***i***_ ⊥ *y*_*i*_ s.t. *p*(*z*_*ij*_ = 1|*y*_*i*_) = *p*(*z*_*ij*_ = 1) = *τ*_*j*,_ and as a result (6) is reduced to

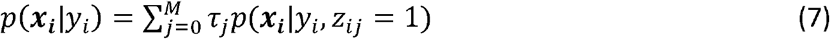

The log likelihood of the observed data adjusting for mutational burden is

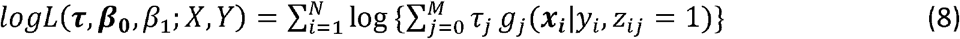

where

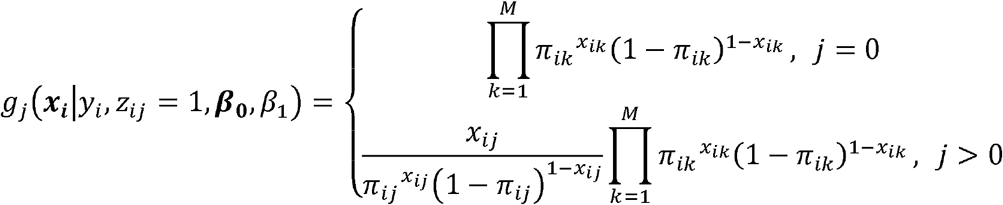

is the cluster-specific probability density function of ***x***_***i***_ conditional on *y*_*i*_, and 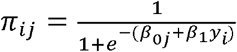. There are no analytical solutions for 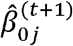 and 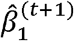in the M-step if using EM algorithm. Thus, we use limited-memory BFGS (L-BFGS)^22^ implemented in PyTorch to minimize the negative log likelihood function and estimate *τ*_*k*_, *β*_0*j*_ and *β*_1_.

### A Statistical Test to Establish Mutual Exclusivity

Our proposed methodology seeks to identify drivers from a framework of observed mutual exclusivity. However, before performing such an analysis on a chosen pathway we propose first conducting a statistical test of the null hypothesis of no mutual exclusivity, i.e., a test of the hypothesis that *γ*=0, or equivalently, ***τ***= (*τ*_0_, *τ*_1_, …, *τ*_*M*_) =(1,0, …,0). In their original development of the MEGSA model, Hua et al. derived an asymptotic likelihood ratio test. Their limiting distribution depends crucially on the assumption that the relative frequencies of driver mutations in each gene are proportional to the passenger mutation rates, an assumption we dropped as indicated earlier. Consequently, we propose to compute the empirical p-value using a parametric bootstrap approach.

Let *θ*=(***τ, β***_**0**_, *β*_1_) denote the parameters in our model. We introduce the following bootstrap estimator for the restricted (under null) and unrestricted settings, respectively.

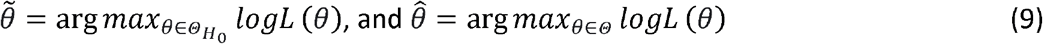

where 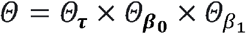 is the full parameter space and 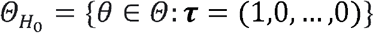.

We propose the bootstrap likelihood ratio statistic

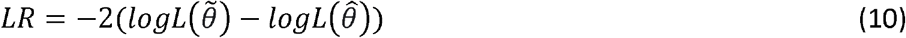

as the test statistic for the null hypothesis.

To construct the test, we generate *B* bootstrap samples (algorithm will be introduced later), denoted by *X*^*(l)*^,*l* = 1,2, …,*B*. Denoting by *LR*^*^ the test statistic from the observed dataset and *LR*^*(l)*^ its value from the *l*^*th*^ bootstrap dataset, the empirical p-value is

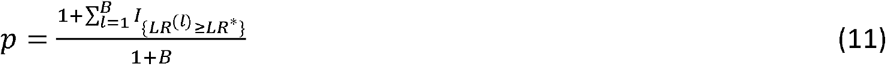

The following data generation algorithm is employed both to create the null distribution (when *γ* = 0) and also to generate datasets under positive levels of mutual exclusivity for our later simulations of model properties. (i) Generate the latent gene membership ***z***_***i***_=(*z*_*i*0_,*z*_*i*1_, …,*z*_*iM*_) of the driver mutation in each tumor *i*. [Note that 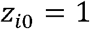, ∀*i* when we are generating a reference distribution under the null hypothesis.] Specifically, 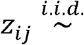 *Multinomial*(1, ***τ***) where ***τ***= (*τ*_0_, *τ*_1_, …, *τ*_*M*_) Tumor *i* has a driver mutation in the *j*^*th*^ gene (*j*=1, …,*M*) if *z*_*ij*_ = 1. Otherwise, it does not possess a driver mutation, and *z*_*i*0_=1. (ii) Generate the centered log-scale mutational burden (*y*_*i*_)for each tumor 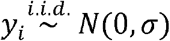. (iii) Generate the individual mutations as *x*_*ij*_ = 1 if *z*_*ij*_ =1 and *x*_*ij*_ ∽*Binomial*(1, *π*_*ij*_)otherwise, where *π*_*ij*_ is computed using equation (5) with given {*β*_0*j*_},*β*_1_ and *y*_*i*_ For simulating data replicates, *σ*,{ *β*_0*j*_} and *β*_1_ are pre-specified. For generating bootstrap data samples, we set 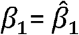, the estimated *β*_1_ by fitting our model to the observed data, and then solve for using *β*_0*j*_ the following equation

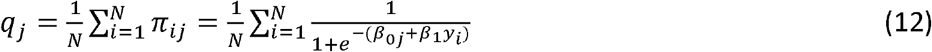

where *q*_*j*_ denotes the overall mutation rate for j^*th*^ gene in the observed data. Equation (12) allows for the bootstrap samples to maintain the association between *π*_*ij*_ and *y*_*i*_ while controlling the similar overall mutation rate for each gene. There is no closed-form solution for *β*_0*j*_ in (12), so we solve for *β*_0*j*_ numerically using Newton’s method.

### Identifying Driver Mutation for Each Tumor and Specific Variants within a Gene

For every tumor we seek to identify the driver mutation, or to determine that there is no driver mutation in the pathway. This can be inferred probabilistically from the posterior probabilities computed through Bayes’ rule. In the absence of adjustment for mutational burden the posterior probability that the mutation in the *j*^*th*^ gene in the pathway is the driver is

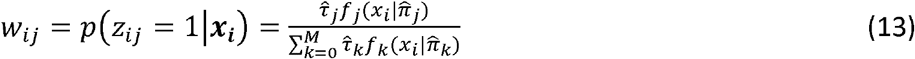

When *j* = 0, equation (13) provides the probability that tumor *i* does not have any driver mutation in the pathway. If a tumor is observed to have mutations in multiple genes, the most likely driver mutation can be determined using

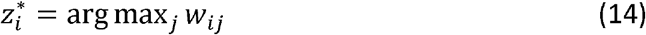

Similarly, the posterior probability under the scenario of adjusting for mutational burden is

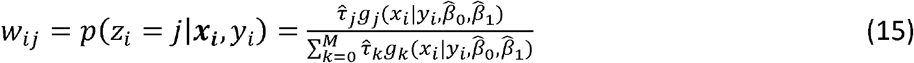

Further, we can gauge the relative influence of individual variants within genes as drivers by averaging these posterior probabilities across the tumors in which the specific variant was observed. Let *x*_*ij(l)*_ denote the mutation status (1-yes; 0-no) of variant *l* from gene *j* in tumor *i*, where variant *l* is nested within gene *j*. The observed mutation frequency for variant *l* (in gene *j*) is

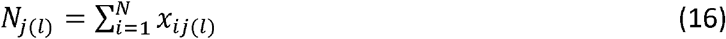

and the average posterior probability that variant *l* is a driver is

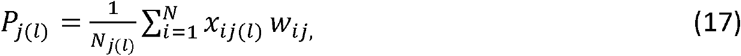

a term we refer to as the “driver frequency”.

## Results

We apply the method to the InterMEL consortium dataset of early-stage melanoma tumors^23, 24^, and then explore, via simulations, the properties of the method.

### Example: Data from the InterMEL Study

The InterMEL study involves genomic sequencing of primary tumors from patients with stage IIA-IIIB melanomas. Our analysis is based on 495 tumor samples genotyped to date. DNA samples were sequenced at Memorial Sloan Kettering Cancer Center using the Integrated <u>M</u>utation <u>P</u>rofiling of <u>A</u>ctionable <u>C</u>ancer <u>T</u>argets or MSK-IMPACT™, a clinically validated and FDA approved hybridization capture-based next-generation sequencing assay developed to guide cancer treatment ^25, 26^. This involved sequencing of 468 cancer-associated genes.

For our illustration of the method, we focus solely on mutations in the RTK-RAS pathway, the major, known pathway that influences the development of melanomas. The MSK-IMPACT panel includes 38 genes from the RTK-RAS pathway. It is well known that mutations occur in melanomas frequently in several genes in this pathway, most prominently the genes *BRAF* and *NRAS*. Mutations in these two genes are almost always mutually exclusive. However, mutual exclusivity has not been studied systematically for other genes in this pathway. Also, for *BRAF* and *NRAS*, hotspot mutations occur very frequently at the 600 locus in *BRAF* and the 61 locus in *NRAS*, but the importance of mutations at other genetic loci in these genes is less clear.

The data reveal that 91% of the 495 tumors had a mutation in the RTK-RAS pathway. However, our estimate of 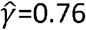 (p-value=0.001) indicates that in only 76% of the tumors is one of the mutations considered to be the driver. Tumor mutational burden varies widely with a standard deviation of 1.1 for the log tumor burden and an estimated effect (on passenger mutation rate) of *β*_1_=1.21. Figure 1 displays data (top panel) and model estimates (bottom panel) for the 18 genes with the highest estimated values of *τ*_*j*_ (proportion of tumors carrying a driver mutation in gene *j*). Figure S1 displays the results for all 38 genes. The top panel displays a structured waterfall plot of the observed mutations. The two most frequently occurring genes at the top of the figure, *BRAF* and *NRAS*, are almost always mutually exclusive, the exceptions being the 4 cases at the extreme left of the figure. It is further noticeable that mutations in these genes frequently occur in the absence of mutations in any of the other genes in the pathway (see the most right-handed columns in the *BRAF* and *NRAS* rows). This high degree of general mutual exclusivity is the key pattern in the data that influences our analysis, confirming a high probability of a driver for mutations observed in these two genes. This is reflected in the bottom panel of Figure 1 where the depth of the color indicates the strength of evidence that the mutation in question is the driver. Quantitative details are provided in Table 1, which displays the relative frequencies for which mutations in the genes occur alongside the portion of these occurrences that are flagged as drivers by of our method. This gene-specific “driver frequency” is the estimated *τ*_*j*_. Interestingly, the method suggests that NRAS is the driver in all tumors involving NRAS mutations, notably the 4 tumors in which *BRAF* and *NRAS* mutations occurred simultaneously. Moving down the gene list the analysis suggests that *KIT* mutations, which occur in about 5% of tumors, is the driver about half of the time, *NF1* mutations are drivers in about ¼ of the 26% of tumors that harbor *NF1* mutations, and that there is limited evidence of driver status for the low frequency genes. Although *NF1* has a relatively high mutation rate, our method downgrades its importance as a driver gene due to its strong association with high tumor mutational burden, which is displayed in the bottom panel of Figure 1, where a dark line indicates that the mutational burden for that tumor is in the top 25^th^ percentile. In general, tumors with high mutational burden are less likely to have a driver mutation identified after model adjustment, but the final posterior probability vector for each tumor computed with the estimated parameter values also depends on other factors (e.g., the proportion of tumors with mutation in a given gene that are singletons). Table S1 summarizes the results for all 38 genes.

**Figure 1.**
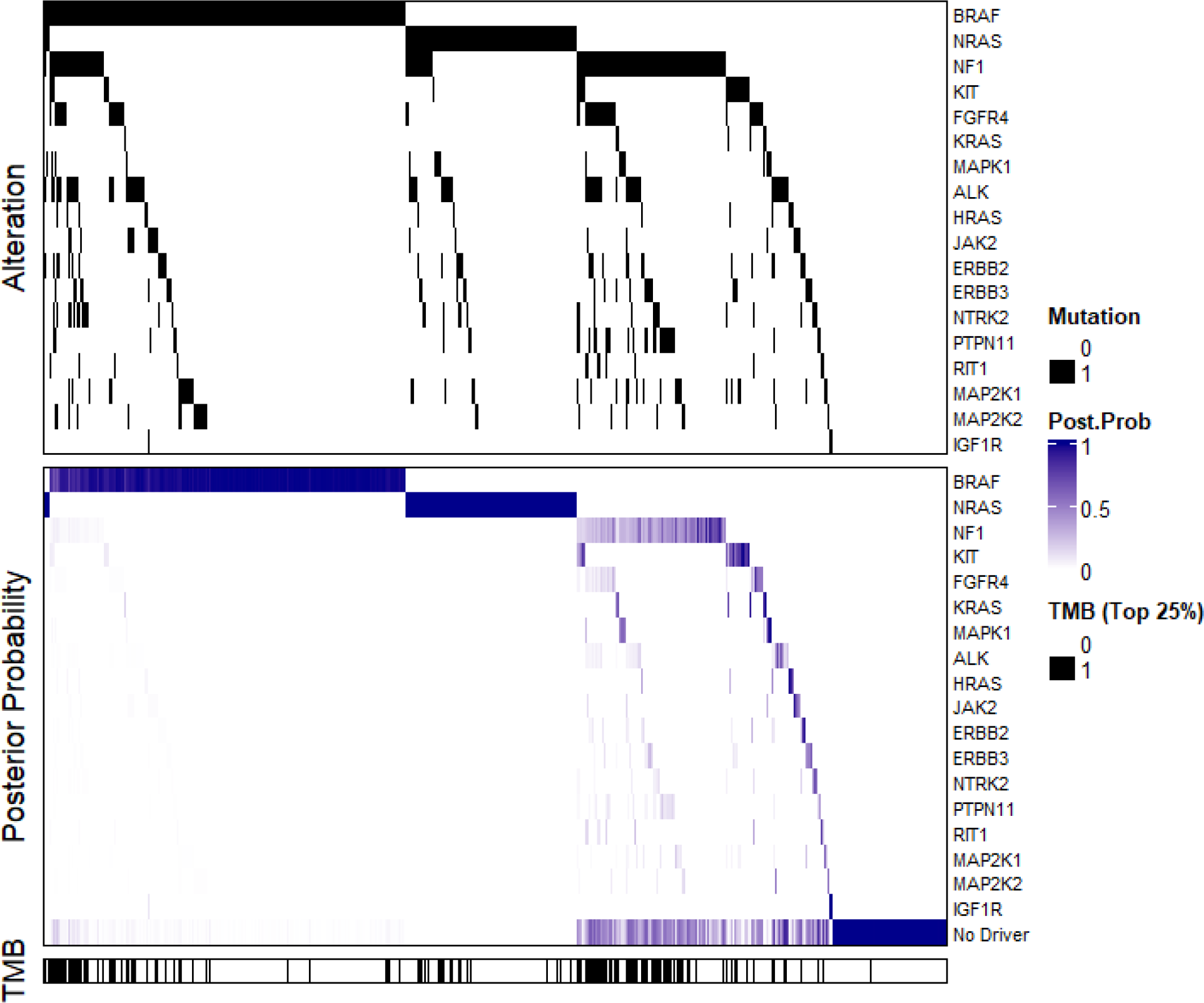
Illustration of the Observed Binary Mutation Status, Estimated Posterior Probability of Driver Mutation, and the Distribution of Binary TMB (Top Genes Only)

**Table 1.**
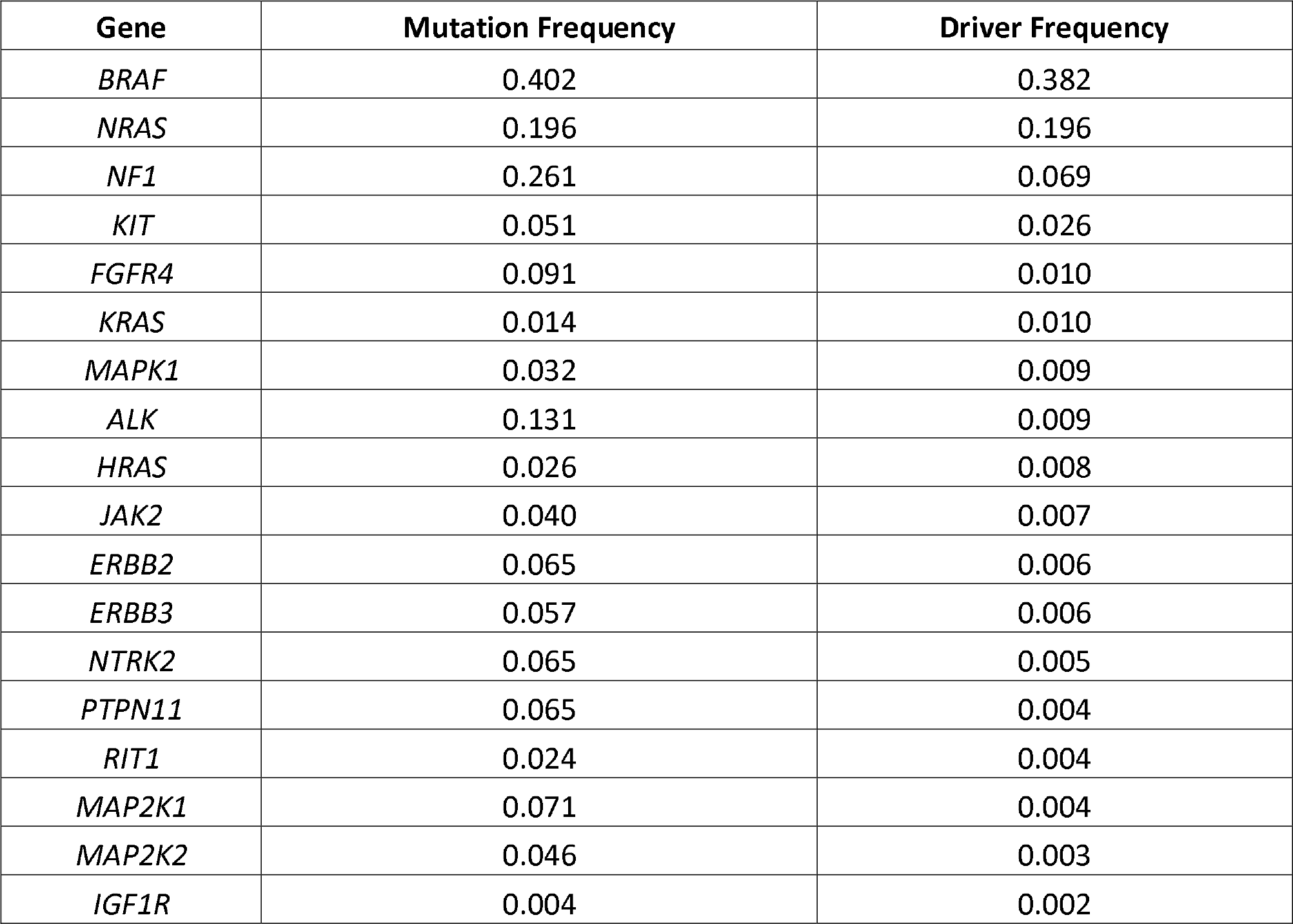
Summary of the Observed Mutation Frequency and the Estimated Driver Frequency for Each Gene in RTK/RAS Pathway (Top Genes Only)

Finally, we focus solely on the *BRAF* gene to illustrate results for individual variants. In Table 2 we provide the frequencies and average posterior probabilities for each of the 48 distinct variants observed. High probabilities are generally assigned when the variant occurs as a singleton, and lower probabilities when other variants in the pathway occur. Very low probabilities occur when a driver variant in a different gene is observed (*NRAS* in this case). Thus the 4 variants at the bottom of this table are the variants (on the extreme left in Figure 1) that occurred alongside an *NRAS* mutation.

**Table 2.**
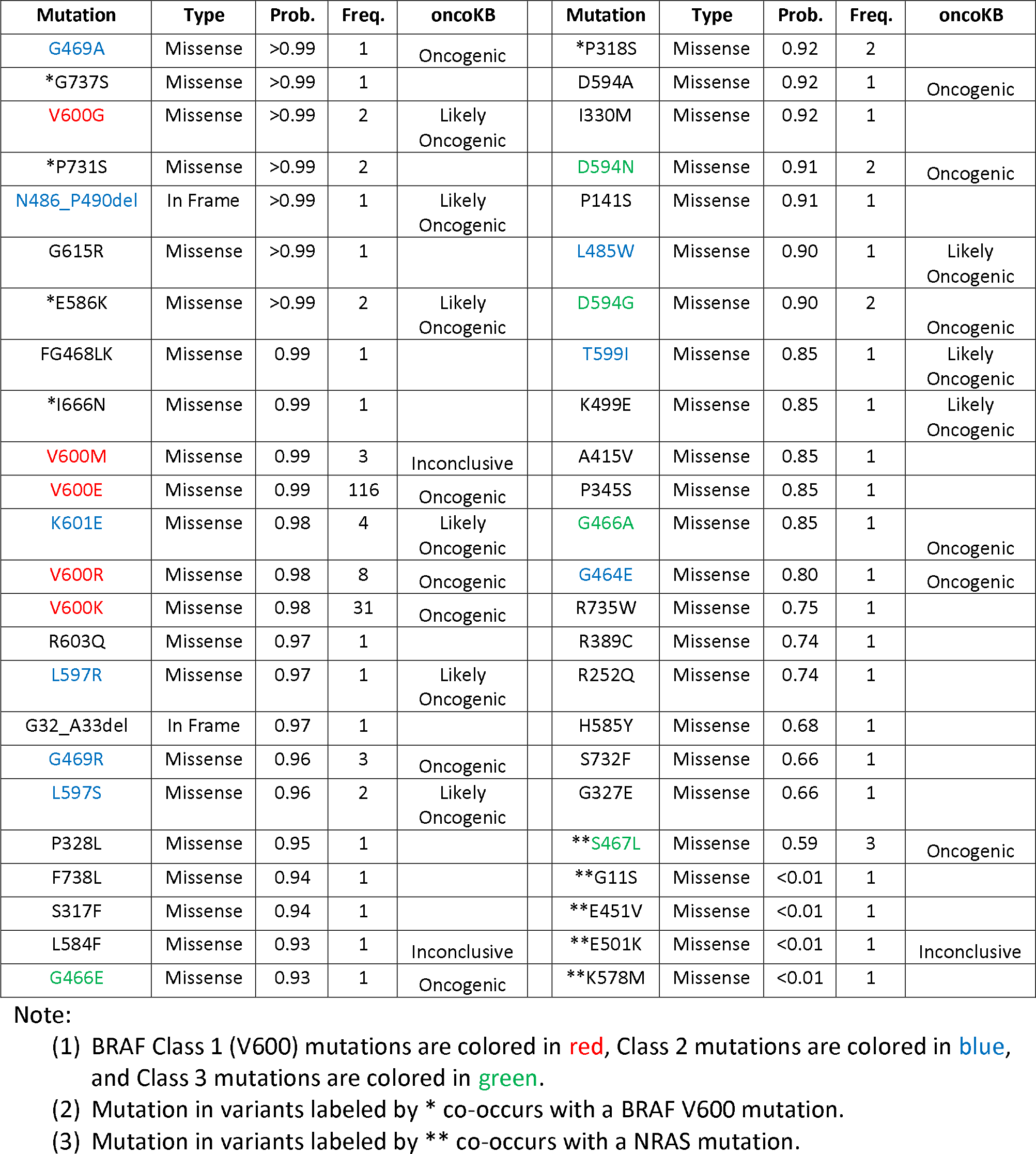
Variant Analysis.

| Mutation | Type | Prob. | Freq. | oncoKB | Mutation | Type | Prob. | Freq. | oncoKB |
| --- | --- | --- | --- | --- | --- | --- | --- | --- | --- |
| G469A | Missense | >0.99 | 1 | Oncogenic | *P318S | Missense | 0.92 | 2 |  |
| *G737S | Missense | >0.99 | 1 |  | D594A | Missense | 0.92 | 1 | Oncogenic |
| V600G | Missense | >0.99 | 2 | Likely<br>Oncogenic | I330M | Missense | 0.92 | 1 |  |
| *P731S | Missense | >0.99 | 2 |  | D594N | Missense | 0.91 | 2 | Oncogenic |
| N486_P490del | In Frame | >0.99 | 1 | Likely<br>Oncogenic | P141S | Missense | 0.91 | 1 |  |
| G615R | Missense | >0.99 | 1 |  | L485W | Missense | 0.90 | 1 | Likely<br>Oncogenic |
| *E586K | Missense | >0.99 | 2 | Likely<br>Oncogenic | D594G | Missense | 0.90 | 2 | Oncogenic |
| FG468LK | Missense | 0.99 | 1 |  | T599I | Missense | 0.85 | 1 | Likely<br>Oncogenic |
| *I666N | Missense | 0.99 | 1 |  | K499E | Missense | 0.85 | 1 | Likely<br>Oncogenic |
| V600M | Missense | 0.99 | 3 | Inconclusive | A415V | Missense | 0.85 | 1 |  |
| V600E | Missense | 0.99 | 116 | Oncogenic | P345S | Missense | 0.85 | 1 |  |
| K601E | Missense | 0.98 | 4 | Likely<br>Oncogenic | G466A | Missense | 0.85 | 1 | Oncogenic |
| V600R | Missense | 0.98 | 8 | Oncogenic | G464E | Missense | 0.80 | 1 | Oncogenic |
| V600K | Missense | 0.98 | 31 | Oncogenic | R735W | Missense | 0.75 | 1 |  |
| R603Q | Missense | 0.97 | 1 |  | R389C | Missense | 0.74 | 1 |  |
| L597R | Missense | 0.97 | 1 | Likely<br>Oncogenic | R252Q | Missense | 0.74 | 1 |  |
| G32_A33del | In Frame | 0.97 | 1 |  | H585Y | Missense | 0.68 | 1 |  |
| G469R | Missense | 0.96 | 3 | Oncogenic | S732F | Missense | 0.66 | 1 |  |
| L597S | Missense | 0.96 | 2 | Likely<br>Oncogenic | G327E | Missense | 0.66 | 1 |  |
| P328L | Missense | 0.95 | 1 |  | **S467L | Missense | 0.59 | 3 | Oncogenic |
| F738L | Missense | 0.94 | 1 |  | **G11S | Missense | <0.01 | 1 |  |
| S317F | Missense | 0.94 | 1 |  | **E451V | Missense | <0.01 | 1 |  |
| L584F | Missense | 0.93 | 1 | Inconclusive | **E501K | Missense | <0.01 | 1 | Inconclusive |
| G466E | Missense | 0.93 | 1 | Oncogenic | **K578M | Missense | <0.01 | 1 |  |

(1) BRAF Class 1 (V600) mutations are colored in red, Class 2 mutations are colored in blue, and Class 3 mutations are colored in green.
(2) Mutation in variants labeled by \* co-occurs with a BRAF V600 mutation.
(3) Mutation in variants labeled by \*\* co-occurs with a NRAS mutation.

*BRAF* variants have been studied extensively and classified into a few major classes with varying potency in oncogenicity based on differences in dimerization requirement and RAS dependency^27, 28, 29, 30^. OncoKB is a widely popular evidence-based variant annotation tool that integrates such known biologic and oncogenic effects^31^. Table 2 shows that the *BRAF* variants with high estimated probability of being a driver (>90%) include most of the functionally potent class I (red) and class II (blue) variants. Also the probabilities are generally higher for variants classified as oncogenic by OncoKB^31^, an important clinical tool for this purpose, providing a partial validation of the method.

### Operating Characteristics of the Method

We have conducted simulations to examine the properties of the method. There are many different features of a pathway that could potentially affect these properties. We have elected to generate data using selected features of the RTK-RAS pathway that were estimated in the previous section, and then to vary some key aspects of these results to explore the influence of selected features. Specifically, we focus on a pathway with 3 types of genes: (1) strong driver genes that function as drivers most of the time (like *BRAF* and *NRAS*); (2) moderate driver genes that sometimes function as drivers and sometimes do not (like *NF1*); and (3) genes that are presumed to never be drivers. Details of their overall and driver frequencies are provided in Table 3. We consider two general configurations, denoted by *A* and *B*. Configuration *A* refers to the low noise setting in which 2 genes are generated from each type. Configuration *B* refers to the high noise setting in which the number of non-driver genes is increased to 10. The probabilities in Table 3 correspond to pathways where mutual exclusivity is present. When we evaluate the test size under the null hypothesis of no exclusivity (Table 4) we use configuration *A* under the assumption that all genes in the configuration have driver rates of 0%. As previously described in the data generation algorithm, the centered log-scale mutational burden was generated from a normal distribution with standard deviation *σ*= 1. *β*_0*j*_ was set to maintain the designed overall and driver mutation frequency under each setting as summarized in Table 3. For all settings we simulated 1000 data replicates under our model structure, and for each test we generated 1000 bootstrap samples.

**Table 3.** Characteristics of Simulated Genes.

| Gene Type | Mutation Frequencies<sup>1</sup> |  | Configuration (# Genes) |  |
| --- | --- | --- | --- | --- |
|  | Overall | Driver | A<br>Low Noise | B<br>High Noise |
| Strong Driver | 20% | 20% | 2 | 2 |
| Moderate Driver | 20% | 10% | 2 | 2 |
| Non-Driver | 20% | 0% | 2 | 10 |

**Table 4.** Size of Test for Mutual Exclusivity at 5% Significance.

| Sample Size | Mutational Burden | Test Size |
| --- | --- | --- |
| 500 | $\beta_1 = 0$ | 0.050 |
| | $\beta_1 = 1$ | 0.046 |
| 1000 | $\beta_1 = 0$ | 0.059 |
| | $\beta_1 = 1$ | 0.055 |
| 5000 | $\beta_1 = 0$ | 0.051 |
| | $\beta_1 = 1$ | 0.047 |

We first examined the properties of the initial significance test to determine the evidence that mutual exclusivity exists in the pathway, using equation (11). We calculated the size of the test for sample sizes ranging from 500 to 5000 under a model in which there was no effect of tumor mutational burden (*β*_1_=0) and under a model in which the effect of tumor mutational burden was in the range of that observed in the real dataset (*β*_1_=0) The results are summarized in Table 4, where the test size is computed as the average proportion of null hypothesis rejections among the 1000 simulated data replicates. We observe that the test size of our proposed bootstrap-based test is in general close to the nominal level of 5% across different settings.

Next, we explored the ability of the model to identify drivers in individual tumors. We used two distinct statistics for this purpose. First, we evaluated overall measures that characterize the true positive and false positive rates for identifying whether or not a tumor has a driver in the pathway. For this calculation the overall false positive rate (FPR) is given by

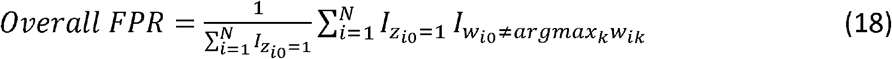

The corresponding true positive rate is given by

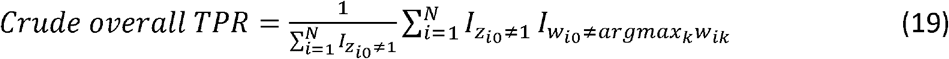

Table 5 summarizes the results of overall accuracy, where FPR and TPR in the table are calculated as the average among the 1000 simulated data replicates. We observe that in general FPR decreases and TPR increases with larger sample size. When there exists an association between passenger mutation rate and tumor mutational burden (i.e., *β*_1_=1) our method tends to classify fewer mutations as drivers, reducing both FPR and TPR. Conversely, elevation in pathway noise tends to make our method nominate more driver mutations, increasing both FPR and TPR. However, the effect of such noise diminishes with larger sample size.

**Table 5.** Overall Accuracy.

| Configuration | Sample Size | Mutational<br>Burden | Accuracy |  |
| --- | --- | --- | --- | --- |
|  |  |  | FPR | TPR |
| **A**<br>**Low Noise** | **500** | $\beta_1 = 0$ | 0.210 | 0.994 |
| | | $\beta_1 = 1$ | 0.156 | 0.968 |
| | **1000** | $\beta_1 = 0$ | 0.211 | 0.999 |
| | | $\beta_1 = 1$ | 0.154 | 0.969 |
| | **5000** | $\beta_1 = 0$ | 0.212 | 1.000 |
| | | $\beta_1 = 1$ | 0.153 | 0.971 |
| **B**<br>**High Noise** | **500** | $\beta_1 = 0$ | 0.247 | 0.997 |
| | | $\beta_1 = 1$ | 0.187 | 0.973 |
| | **1000** | $\beta_1 = 0$ | 0.213 | 0.999 |
| | | $\beta_1 = 1$ | 0.166 | 0.973 |
| | **5000** | $\beta_1 = 0$ | 0.212 | 1.000 |
| | | $\beta_1 = 1$ | 0.155 | 0.972 |

Finally, we explored diagnostic accuracy at a more granular level, seeking to determine the accuracy of driver identification for the different individual gene configurations. For this purpose, we define the gene-specific false positive and true positive rates. That is, our false positive rate in this context measures, among tumors with a mutation in gene *j* that is not a driver, what proportion are incorrectly flagged as a driver

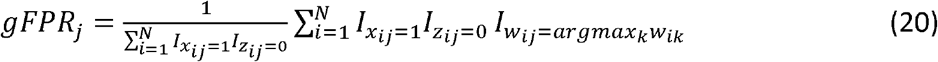

The corresponding true gene-specific positive rate measures, among tumors with a mutation in gene *j* that is a driver, what proportion are correctly flagged as a driver

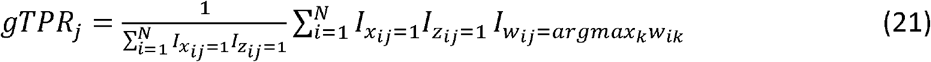

Results are provided in Table 6, where, as before, the gFPR and gTPR values are averages across all simulated datasets. It is worth noting that the previously defined overall FPR and crude overall TPR are not simple weighted averages of gFPR or gTPR across genes as the denominators are in those formulas are not the same. We do not compute gFPR for strong driver genes because in our simplified construct there are no tumors which have non-driver mutations in these genes (i.e., no passenger mutations identified). Similarly, we do not compute gTPR for non-driver genes. For strong driver genes, we observe that the average gTPR is close to 1 in almost all scenarios (e.g., small sample size or high noise). Similarly, the method also performs well in screening out non-driver genes, evidenced by the extremely small gFPR across different scenarios. For moderate driver genes, those that can often be either a driver or a passenger, it is clearly more challenging for the method to identify drivers accurately, with gFPRs ranging from 0.340 to 0.477 across our various configurations, while the gTPRs range from 0.851 to 0.942. As was shown previously in Table 5, with greater association between passenger mutation rate and tumor mutational burden our method tends to classify fewer mutations as drivers, reducing both FPR and TPR. Conversely, the presence of increasing noise has minimal impact at an individual gene level.

**Table 6.** Gene-Specific Diagnostic Accuracy Rates.

| Configuration | Sample Size | Mutational<br>Burden | Gene Type | Accuracy |  |
| --- | --- | --- | --- | --- | --- |
|  |  |  |  | gFPR | gTPR |
| **A**<br>**Low Noise** | **500** | $\beta_1 = 0$ | Strong Driver | | 1.000 |
|  |  |  | Moderate Driver | 0.471 | 0.920 |
|  |  |  | Non-Driver | 0.001 |  |
| | | $\beta_1 = 1$ | Strong Driver | | 0.999 |
|  |  |  | Moderate Driver | 0.342 | 0.851 |
|  |  |  | Non-Driver | 0.004 |  |
| | **1000** | $\beta_1 = 0$ | Strong Driver | | 0.999 |
|  |  |  | Moderate Driver | 0.475 | 0.937 |
|  |  |  | Non-Driver | <0.001 |  |
| | | $\beta_1 = 1$ | Strong Driver | | 0.999 |
|  |  |  | Moderate Driver | 0.343 | 0.859 |
|  |  |  | Non-Driver | 0.001 |  |
| | **5000** | $\beta_1 = 0$ | Strong Driver | | 0.999 |
|  |  |  | Moderate Driver | 0.476 | 0.942 |
|  |  |  | Non-Driver | <0.001 |  |
| | | $\beta_1 = 1$ | Strong Driver | | 0.999 |
|  |  |  | Moderate Driver | 0.340 | 0.867 |
|  |  |  | Non-Driver | <0.001 |  |
| **B**<br>**High Noise** | **500** | $\beta_1 = 0$ | Strong Driver | | 1.000 |
|  |  |  | Moderate Driver | 0.475 | 0.931 |
|  |  |  | Non-Driver | 0.008 |  |
| | | $\beta_1 = 1$ | Strong Driver | | 0.999 |
|  |  |  | Moderate Driver | 0.362 | 0.867 |
|  |  |  | Non-Driver | 0.005 |  |
| | **1000** | $\beta_1 = 0$ | Strong Driver | | 0.999 |
|  |  |  | Moderate Driver | 0.477 | 0.940 |
|  |  |  | Non-Driver | <0.001 |  |
| | | $\beta_1 = 1$ | Strong Driver | | 0.999 |
|  |  |  | Moderate Driver | 0.352 | 0.869 |
|  |  |  | Non-Driver | 0.002 |  |
| | **5000** | $\beta_1 = 0$ | Strong Driver | | 1.000 |
|  |  |  | Moderate Driver | 0.475 | 0.942 |
|  |  |  | Non-Driver | <0.001 |  |
| | | $\beta_1 = 1$ | Strong Driver | | 0.999 |
|  |  |  | Moderate Driver | 0.344 | 0.870 |
|  |  |  | Non-Driver | <0.001 |  |

## Discussion

Our goals in developing this methodology were to find a strategy for identifying potential driver mutations in a tumor and assigning probabilities to the potential candidates. We built our strategy on a model that frames the selection on the presence of mutual exclusivity patterns in the data. Among the many groups that have studied mutual exclusivity in this context, we elected to build on the ideas of Hua et al.^15^ since their model was firmly based on well-established statistical principles. The underlying model is structured around the assumption that there can be at most 1 driver in the pathway in any individual tumor, and this is in itself an assumption that may not be correct. However, this assumption does provide a solid framework in which to examine mutual exclusivity and, as we have seen with the RTK-RAS example, produces results that appear to be highly plausible in that they align with known evidence about this pathway. However, the RTK-RAS example, though useful for illustration, represents a pathway for which the mutual exclusivity between *BRAF* and *NRAS* is especially profound, and thus may be an easier task for our model than pathways without highly prevalent variants that are very strongly mutually exclusive.

We believe that our method has strong potential for shedding light on which specific mutations are potentially pathogenic in a specific gene. In the *BRAF* example we presented, the V600 variants identified as pathogenic are well characterized and are targets for FDA approved therapies. However, approximately 35% of all *BRAF* mutations occur outside the V600 codon^29^. The functional impact and therapeutic potential of non-V600 *BRAF* mutations is an active research topic, yet existing knowledge in this area is limited. Our analysis of *BRAF* identified variants other than the common V600 variants that may be potentially pathogenic. These represent the kinds of variants that could be prime candidates for experimental validation using modern in vitro and in vivo strategies^32^.

We emphasize that our strategy is focused on a single pathway and based on the pivotal assumption that there can be only one driver in the pathway in any given tumor. However, in any given tumor there are very likely multiple drivers, each occurring in distinct pathways. While one could perform our analysis independently for distinct pathways in order to identify a more complete set of drivers, a future research task is to expand our approach to permit a simultaneous analysis of multiple pathways.

## Acknowledgments

This study was supported by the National Institutes of Health (P01 CA206980 and R01 CA251339) and Memorial Sloan Kettering Cancer Center (P30 CA008748).

## Code availability

Our algorithm is implemented in Python and available on GitHub at https://github.com/tarot0410/MAGPIE.

## Notes

### Competing Interest Statement

The authors have declared no competing interest.

