## Supplemental Figure 1 and Supplemental Table 1 for "Adaptation of a Mutual Exclusivity Framework to Identify Driver Mutations within Biological Pathways"

**[1]Department of Epidemiology and Biostatistics, Memorial Sloan Kettering Cancer Center, New York, NY, USA**

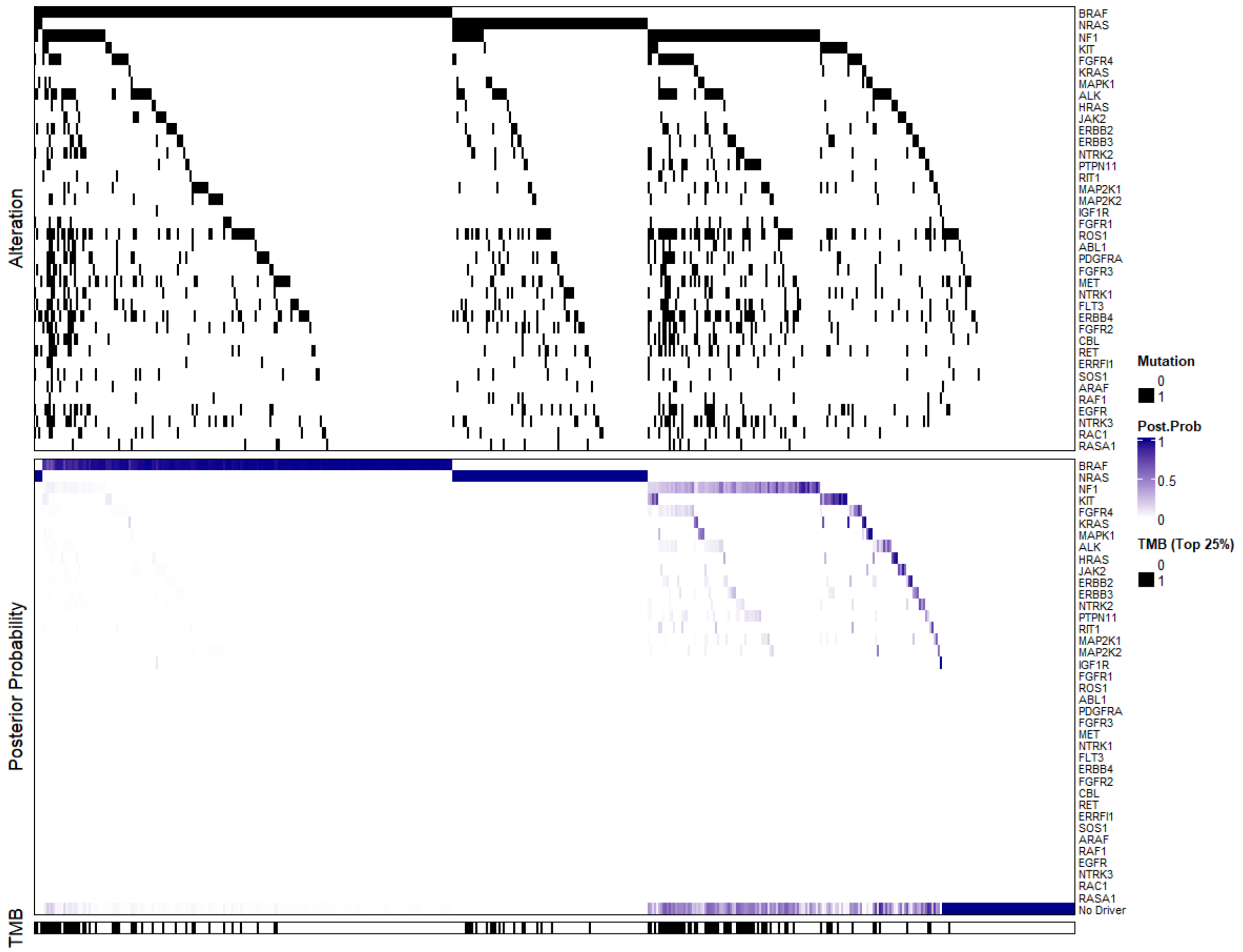


**Figure S1: Illustration of the Observed Binary Mutation Status, Estimated Posterior Probability of Driver Mutation, and the Distribution of Binary TMB (All 38 Genes).**

**Table S1. Summary of the Observed Mutation Frequency and the Estimated Driver Frequency for Each Gene in RTK/RAS Pathway (All 38 Genes)**

| Gene | Mutation Frequency | Driver Frequency |
|---|---|---|
| *BRAF* | 0.402 | 0.382 |
| *NRAS* | 0.196 | 0.196 |
| *NF1* | 0.261 | 0.069 |
| *KIT* | 0.051 | 0.026 |
| *FGFR4* | 0.091 | 0.010 |
| *KRAS* | 0.014 | 0.010 |
| *MAPK1* | 0.032 | 0.009 |
| *ALK* | 0.131 | 0.009 |
| *HRAS* | 0.026 | 0.008 |
| *JAK2* | 0.040 | 0.007 |
| *ERBB2* | 0.065 | 0.006 |
| *ERBB3* | 0.057 | 0.006 |
| *NTRK2* | 0.065 | 0.005 |
| *PTPN11* | 0.065 | 0.004 |
| *RIT1* | 0.024 | 0.004 |
| *MAP2K1* | 0.071 | 0.004 |
| *MAP2K2* | 0.046 | 0.003 |
| *IGF1R* | 0.004 | 0.002 |
| *FGFR1* | 0.038 | <0.001 |
| *ROS1* | 0.242 | <0.001 |
| *ABL1* | 0.061 | <0.001 |
| *PDGFRA* | 0.103 | <0.001 |
| *FGFR3* | 0.073 | <0.001 |
| *MET* | 0.141 | <0.001 |
| *NTRK1* | 0.091 | <0.001 |
| *FLT3* | 0.097 | <0.001 |
| *ERBB4* | 0.186 | <0.001 |
| *FGFR2* | 0.111 | <0.001 |
| *CBL* | 0.071 | <0.001 |
| *RET* | 0.097 | <0.001 |
| *ERRFI1* | 0.028 | <0.001 |
| *SOS1* | 0.051 | <0.001 |
| *ARAF* | 0.024 | <0.001 |
| *RAF1* | 0.030 | <0.001 |
| *EGFR* | 0.101 | <0.001 |
| *NTRK3* | 0.127 | <0.001 |
| *RAC1* | 0.073 | <0.001 |
| *RASA1* | 0.038 | <0.001 |
